# A major entomoparasite interferes with Chikungunya transmission by *Aedes albopictus*

**DOI:** 10.1101/2024.12.04.626309

**Authors:** Edwige Martin, An-nah Chanfi, Barbara Viginier, Vincent Raquin, Claire Valiente Moro, Guillaume Minard

## Abstract

The Asian tiger mosquito, *Aedes albopictus*, spreads diseases like Chikungunya and has caused outbreaks worldwide. Studies show that mosquitoes-associated microbes can affect how well they spread diseases. One of those microbes, the parasite *As. taiwanensis*, is common in older mosquito populations but rare in newer ones. We found that this parasite slows down the spread of Chikungunya within the mosquito and decreases its transmission rate by half. Unparasitized mosquitoes spread the virus more easily, suggesting that changes in mosquito microbes could impact disease outbreaks and public health.

---

The Chikungunya virus is an arbovirus (*i.e*. arthropod borne virus) that causes febrile illness, a rash as well as severe and often debilitating arthralgias that may last for several years in humans. Originally, it was mostly vectored by the Yellow fever mosquito *Aedes aegypti* but, due to a single mutation (*i.e*. A226V), its vector specificity has shifted toward a higher transmission by the Asian tiger mosquito *Ae. albopictus* (1).

Originating from southern and eastern Asia, *Ae. albopictus* is the most invasive mosquito species and has recently been introduced in every inhabited continents due to global trade of products containing mosquito eggs (2). The combined impact of mosquito invasiveness and circulation of CHIKV strains carrying the A226V mutation has drastically favored the emergence of explosive chikungunya outbreaks. The Indian Ocean, in particular, was strongly impacted between 2004 and 2007 with one third of the population from La Réunion island being infected and 10% of the patients presenting disabling after-effects 13 years later (3). To be transmitted from one individual to another the virus need to be acquired by a competent female mosquito when biting an infected host (4). Following this natural entry route, the virus replicates into the mosquito midgut epithelium to get disseminated throughout the mosquito body and reach the salivary glands. CHIKV particles are then transmitted to a healthy host through saliva that is secreted during a blood meal.

During the last decades, many studies have suggested that pathogen transmission can be modulated by mosquito associated microorganisms forming their microbiota (5). Some microorganisms interfere with the mosquito vector competence that is defined by their ability to efficiently get infected, replicate and retransmit a given pathogen. The microbiota of a given mosquito can (i) directly interfere with pathogens for example by lysing them (6) or (ii) indirectly interfere with pathogens by reinforcing the insect’s natural barrier such as their innate immunity (7). However, we are still lacking information on the consequences of variations in the mosquito microbiota toward pathogens transmission. *Ae. albopictus* prokaryotic and eukaryotic microbiota has been reported to vary among populations (8). If all the populations and almost all the individuals are infected by the dominant intracellular bacteria *Wolbachia w*AlbA and *w*AlbB they show marked variations in their association with the second most dominant microorganism, the extracellular Apicomplexan protist *Ascogregarina taiwanensis*.

Indeed, *As. taiwanensis* colonizes more than 80% of individuals in native or old mosquito populations while it colonizes less than 20% of the individuals in recently introduced ones (*i.e*. up to 2 years after introduction) presumably due to its weak ability to survive mosquito egg transportation conditions (8). This protist is considered as a weak mosquito parasite colonizing the adult Malpighian tubules and that is only costly for its host when conditions are harsh (9). Preliminary studies evidenced that the presence of *As. taiwanensis* reduces the density of the pathogenic dog earthworm *Dirofilaria immitis* in *Ae. albopictus* (10). In this study we propose to investigate its effect on *Ae. albopictus* vector competence toward CHIKV. To do so, we compared the dynamic of infection, dissemination and transmission of the virus over the time after unparasitized and parasitized mosquitoes have been fed with an infectious blood meal (**Figure 1A**). The experiment was repeated twice on two mosquitoes’ generations. We have then correlated the densities of viral particles and parasites (*i.e*. oocysts form) in coinfected individuals.

**Figure 1.**
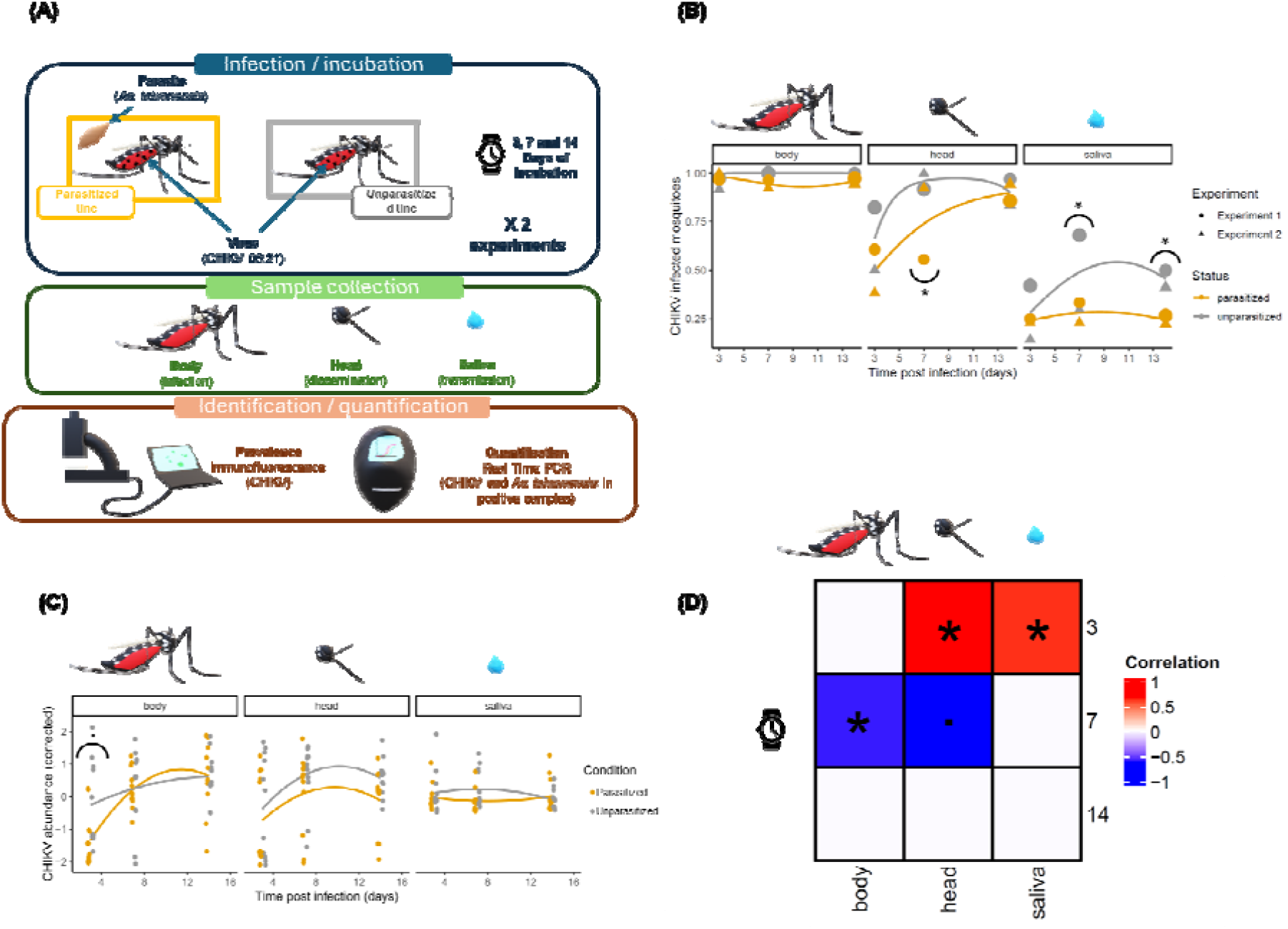
Impact of the parasite on *Ae. albopictus* vector competence toward CHIKV. (**A**) Parasitized and unparasitized mosquitoes were exposed to a blood meal infected with the CHIKV 06.21 (∼10^7^ FFU.ml^-1^) strain. The experiment was repeated twice on different mosquito generations. A total of 10 to 38 engorged females were collected for each experiment and each time point after an incubation of 3-, 7-or 14-days post blood meal. The body, head and saliva of mosquitoes were collected, and detection of the virus was performed for each sample by immunofluorescence on C6/36 cell lines. A total of 5-6 samples in which the virus was detected were selected for real time PCR quantification of the virus and the parasite. (**B**) For each sample type and each time point (in days post infection) two experiments were performed. Dots represent CHIKV prevalence (infection rate) values for each combination of sample types, time point and experiment. The dot size is proportional to the sample size in each essay (from 10 to 38). Lines represent a second order polynomial regression of prevalence over time. Mosquitoes that are parasitized by *As. taiwanensis* are represented in orange while mosquitoes that are unparasitized are represented in grey. (**C**) For each sample type and each time point, relative CHIKV abundances were represented after correcting variations in quantifications that can be attributed to the experiment. Lines represent a third order polynomial regression of quantifications over the time. Mosquitoes that are parasitized by *As. taiwanensis* are represented in orange while mosquitoes that are unparasitized are represented in grey. (**D**) For mosquitoes that were co-infected by the parasite and the virus, correlations were performed between abundance sof *As. taiwanensis* and CHIKV particles for each time point (in rows) and each sample type (in columns). Positive spearman correlations are represented in blue, while negative correlations are represented in red. For each graph p-value results from Tukey-HSD *post-hoc* tests (**B, C**) or Spearman correlations (**D**) were highlighted. Significant correlations (*) or tendency (.) were both considered *i.e*. p-value < 0.05 or <0.1 respectively.

Prevalence of the virus was estimated by immunofluorescence on C6/36 cells in body, head or saliva samples form parasitized and unparasitized mosquitoes 3, 7 and 14 days after being fed with an infectious blood meal (*i.e*. days post infection – dpi). Infection rate was measured as the proportion of positive mosquito bodies for the virus presence. It reached almost 100% and it was not influenced by the mosquito parasitic status and was stable over time (**Figure 1B; Table S1**). Conversely, the proportion of mosquitoes that exhibit virus dissemination in head increased over the time regardless of their parasitic status of mosquitoes (**Table S1**). However, dissemination of the virus to the head was slightly delayed in parasitized mosquitoes. Indeed, at 7 dpi, unparasitized mosquitoes were 6.74±4.63 times more likely to harbor disseminated viruses in mosquito heads than their parasitized conspecifics (**Figure 1B**). Although the difference was not significant at other time points with on average 64±0.05% of infected mosquito heads at 3dpi and 90.2±0.03% at 14dpi. The proportion of females being able to transmit CHIKV *via* saliva did not significantly differ over the time (probably due to a lack of statistical power), while it was significantly influenced by the parasitic status of mosquitoes (**Table S1**). In more details, the proportion of mosquitoes that released CHIKV *via* saliva at 3 dpi was relatively homogenous between both parasitic status, while it was lower for parasitized individuals with 3.13±1.6 and 2.66±1.19 time less chances to transmit viral particles at 7 and 14 dpi (**Figure 1B**).

Viral RNA load was estimated by RT-qPCR of the E2 virus gene and normalized by the mosquito RPS17 housekeeping gene for 5 to 6 samples that were previously identified as infected by the virus and for each experiment, each time point and each sample. The abundance of viral particles increased over time in the body and head of mosquitoes, regardless of their parasitic status type (**Figure 1C**; **Table S1**). A slight tendency was also observed toward an interaction between the mosquito condition and time post infection. Indeed, CHIKV abundances were slightly lower in the body of parasitized mosquitoes compared to their unparasitized relatives at early time point while no tendency was observed in the later ones. No differences were observed in CHIKV abundances between parasitized and unparasitized mosquitoes in the head and saliva (**Figure 1C**; **Table S1**).

When individuals were simultaneously co-infected by the parasite and the virus, we also checked whether the parasite density correlate with the density of viral particles that infect the mosquito, disseminate into its head or are transmitted through saliva over the time. At 3 dpi, abundances of parasite were positively correlated with abundances of viral particles in the mosquito head and saliva (**Figure 1D; Table S2)**. This suggests that when mosquitoes are coinfected with CHIKV and the parasite, individuals that harbor higher parasite densities are most likely to be associated with higher abundances of viral particles that disseminate into the head and are transmitted by the mosquito for early time points while their body is infected by the same density of CHIKV, reflecting higher virus dissemination efficiencies. Interestingly, at 7 dpi the trend tends to differ. The parasite abundance is negatively correlated with the viral particles’ abundances in the mosquito body and head while no significant correlation was observed in the mosquito saliva. This suggests that infected individuals harboring higher parasite densities tend to be infected by a lower density of viral particles that poorly disseminate. Finally, no correlation was observed between the parasite and CHIKV abundances in the body, head and saliva of the mosquito at 14 dpi.

In this study we provide the first proof of concept results showing that *Ascogregarina* parasites alter the vector competence of *Aedes albopictus* toward CHIKV. To do that we used a mosquito population for which we previously estimated CHIKV 06.21 intra-vector dynamics for a range of human viremia-like CHIKV doses (4). The dynamic reported in that former study was very close to that of the non-parasitized individuals in this current investigation. Therefore, deviation of dissemination and transmission in parasitized mosquitoes can be confidently inferred to the interaction of individuals with *As. taiwanensis*. Until now, *Wolbachia* is the mosquito-associated major microbial symbiont for which viral interference was intensively studied (11,12). Mechanisms behind its antiviral activities remain elusive but interactions with lipid metabolism (in particular cholesterol scavenging) (13), cellular autophagy (14), oxidative stress and innate immunity (15) have been proposed.

*As. taiwanensis*-mediated modulation of the insect immune system has previously been observed (10) and a genome survey of the parasite suggests that it could contribute to the lipid metabolism of its host (16). We noted that the parasite density was negatively correlated with the virus density only 7 days after infection. Therefore, the interaction may need some latency as it is often observed for interactions that require activations of a pathway (*i.e*. such as immunity or metabolism). Direct interactions with the virus may exist during the adult stage. However, the virus and the parasite are not colocalized in key organs for virus replication since the parasite colonizes Malpighian tubules of the mosquito (17) that are posterior to the midgut where CHIKV replicates before its dissemination to salivary glands through the hemolymph. It is important to note that *As. taiwanensis* colonizes the epithelial cells of the mosquito midgut during the 2^nd^ – 4^th^ larval instars before being released in the gut lumen and colonizing Malpighian tubules during pupation (17). This may induce long lasting fragilization of the gut structure that may in turn facilitate virus entry into the mosquito hemocoel (*i.e*. body cavity containing the hemolymph) and facilitate its dissemination. The midgut barrier is indeed a major bottlenecks that filters out most of the virions (18).

Previous studies showed that the microbiota of *Ae. albopictus* is less diverse in recently introduces populations compares to native ones (8,19). This is partly due to the loss of this parasite, a dominant member of the native population’s microbiota, which is lost during the transportation of mosquito eggs. As a result, recently introduced populations experience a parasite release lasting nearly two years. Considering our knowledge on the distribution of this parasite we suggest that its negative impact on mosquito vector competence could be considered in the near future as a proxy for improving epidemiological risk estimation. However, other factors that lead to variations in the transmission of CHIKV (*e.g*. other members of the microbiota, vector biting rate, survival, and density) should also be considered and integrated in such models (5,20). Further, experimental studies are also mandatory to determine whether those factors interact with *As. taiwanensis* influence on vector competence of *Ae. albopictus* toward CHIKV in the field.

In conclusion, we provide the first evidence on the impact of a natural and prevalent member of the *Ae. albopictus* microbiota on one of the major arboviruses it transmits. The interference effect of the Apicomplexa *As. taiwanensis* against *Ae. albopictus* is a promising step toward the understanding of microbiota-induced disparities in mosquitoes vector competence. We evidenced that parasitism of the mosquito delays the dissemination and decreases the transmission of CHIKV particles by infected mosquitoes. Since recently introduced mosquitoes’ populations escaped from the parasite compared to old and native ones, those results may help to better model the epidemiological risk related to CHIKV transmission over the time after *Ae. albopictus* invade a new territory. However, further studies of *As. taiwanensis* on the mosquito fitness and behavior as well as its interaction with the rest of the microbiota and their underlying consequences for virus transmission may be required to better estimate the CHIKV transmission risk. Finally, a better understanding of those interactions may contribute to the development of biocontrol strategies based on the utilization of such microbial agents as already experimented in the field for the *Wolbachia* endosymbiont.

## Acknowledgement

This study was funded by the ANSES project PNR-EST 2019 AS-CONTROL as well as the ANR JCJC Project Evasion. We thanks the FR BioEEnViS for providing us access to the Molecular Biology DTAMB platforms. We acknowledge the contribution of SFR Biosciences (UAR3444/CNRS, US8/Inserm, ENS de Lyon, UCBL) AniRA-L3 facility (Marie-Pierre Confort) for their help. The traineeship of AC was performed within the cursus of the Master “Microbiologie” from Clermont-Ferrand.

## Ethics statement

Parasitized and unparasitized female mosquitoes were fed on anesthetized mice during the amplification steps of the populations. This protocol was reviewed by the Institutional Animal Care and Use Committee, acceptance reference number: Apafis #31807-2021052715018315. Use of human blood donors for the mosquito infectious blood meal was enabled by an agreement with the French National Blood Bank: CODECOH DC-2019-3507.

## Supporting information

### Materials and methods

#### Mosquito lines

Parasitized and unparasitized mosquito strains were generated from a mosquito population that has been collected in Villeurbanne and Pierre-Bénite in 2017 and presenting natural infection of *Ascogregarina taiwanensis*. Further information on mosquito strains generation and maintenance were previously detailed (21). We verified the presence and absence of the parasite in each strain by microscopy and diagnostic qPCR (see protocol details in the next sections).

#### Mosquito exposure to Chikungunya virus by artificial blood meal

Experiments were conducted in level 3 biosafety facility (BSL3). Mosquitoes were exposed to CHIKV 06.21 (provided by Anna-Bella Failloux, Paris Pasteur Institute) originating from La Réunion Outbreak (2005-2006) and carrying the A226V mutation. Virus stock was produced on Vero E6 cells at a multiplicity of infection of 0.01 in DMEM medium (Merck, Germany) supplemented with 10% of fetal bovine serum (Cytiva, USA). After 72h of incubation at 37°C, 5%CO_2_, the medium was centrifuged at 200g for 5min at 4°C to remove cellular debris. Supernatant was harvested, mixed with 5.750ml of Hepes (50mM) sucrose (500mM) solution and stored at –80°C as aliquots. Stock infectious titers were estimated by fluorescent focus assay on C6/36 cells (derived from *Ae. albopictus* larvae) (4). The day before infectious blood meal, fresh human blood (originating from multiple human anonymous donors, French National Blood Bank – Etablissement Français du Sang) was washed 3 times in PBS to recover erythrocytes. Mosquitoes were placed in feeding boxes (*i.e*. plastic containers covered with a net) then transferred at 26°C in the BSL3. The day of the infectious blood meal, CHIKV stock was mixed with the erythrocytes suspension (1:2, v:v) to reach an infectious titer of 10^7^ FFU.ml^-1^ in the blood meal. Blood feeding was performed with Hemotek feeders (Hemotek) covered with porc intestine and filled with 2.5mL of infectious blood heated at 37°C. Female mosquitoes were allowed to feed for 25 min. After blood meal, they were anesthetized, on ice, at 4°C for 30 min and fully engorged females were transferred in 1-pint cardboard cup (up to 25 females/cup) then incubated at 26.5°C, 16h/8h light/dark and 80% of humidity in presence of a cotton imbibed with a 10% sucrose solution. Aliquots of infectious blood were stored –80°C to control CHIKV infectious titer upon blood meal duration. Two independent blood meals were performed with two generation of parasitized and unparasitized females. The measured CHIKV titer measured at the end of each blood meal was 0.93×10^7^ FFU.ml^-1^ (experiment 1) and 1.15×10^7^ FFU.ml^-1^ (experiment 2).

#### Vector competence assay

Female mosquitoes were collected at 3, 7 or 14 days post infectious blood meal and processed to measure infection, dissemination and transmission rates by estimating CHIKV prevalence in mosquito body, head and saliva respectively. For each time point, individuals were anesthetized on ice. Their legs and wings were removed and they were glued on a plate with double sided tape (3M). Their proboscis was inserted in a 10µL filter tip containing 10µL of Fetal Bovine Serum (Cytiva) to collect saliva, as described (22). A 1.5 µL drop of 1% pilocarpine (Merck) supplemented with 0.1% Tween-20 (Sigma) was added on the thorax of each female to stimulate salivation in the filter tip. Salivations were performed for 1h at 26.5°C. Saliva solution transferred in a tube containing 100µL of culture medium (DMEM 1X with 4% FBS, 2.5µg/mL of Amphotericin, 100U/mL of Nystatin, 50µg/mL of Gentamicin, 100U/ml of Penicillin, 100µg/mL Streptomycin) and stored at –80°C. Heads and bodies of each individual were harvested and transferred in tubes containing 400µL of culture medium and a 3mm tungsten bead (Qiagen). Those tissues were crushed by agitation in a Qiagen Tissue Lyser II for 1min at 30Hz and stored at –80°C.

#### Estimation of CHIKV prevalence

CHIKV detection was performed on 96 well plates coated containing 3×10^5^ C6/36 cells and supplemented with 40µL of saliva, head or body samples. Plates were incubated for 3 days at 28°C. They were fixed with 150µL of 4% paraformaldehyde (Santa Cruz Biotechnology) for 20 min at room temperature. The overlay was removed and cells were washed three times with 100µL of DPBS (ThermoFisher scientific). Cell permeabilization was conducted by adding 50µL of a DPBS solution supplemented with 0.3% Triton-X 100 (Sigma) and 1% of Bovine Serum Albumin (BSA, Cytiva) for 30min at 37°C. Samples were washed three times with 100µL of DPBS. A volume of 50µL of a Semliki Forest virus anti-capsid antibody cross-reacting with CHIKV diluted 1:600 in DPBS and 1% BSA was added in each well for 1h at 37°C. This antibody was previously shown to be sensitive toward CHIKV (23). Cells were washed three time with 100µL of DPBS before being incubated for 1h at 37°C with 50µL of a 1:500 diluted goat anti-mouse secondary antibody coupled with an Alexa fluor 488 fluorochrome (Invitrogen). They were then washed three more times with DPBS before being observed under an epifluorescence microscope (Zeiss Colibri 7) at a 100X magnification. Samples for which the cell layers contained fluorescent foci were scored positive for CHIKV while those for which no fluorescent foci were observed were considered as uninfected. Positive (CHIKV stock) and negative (culture media) controls were performed on each titration plate. Infection, dissemination and transmission efficiencies corresponded to the prevalence of CHIKV (*i.e*. proportion of positive samples) in body, head and saliva respectively.

#### Quantification of CHIKV and *Ascogregarina taiwanensis*

Total RNA isolation was performed using 30µL of mosquito sample (either saliva, head or body) mixed with 70µL of TRIzol (Invitrogen). After 10 min at room temperature, 20µL of chloroform were added then vortexed and centrifuged for 15min at 17,000g, 4°C. The supernatant was collected and mixed with 60µL of isopropanol supplemented with 1µL of glycoblue (Invitrogen) before being homogenized and stored overnight at –20°C. Samples were then centrifuged for 15min at 17,000g and 4°C. Supernatant was discarded and the pellet was resuspended in 500µL of 70% cold ethanol before being pelleted by centrifugation for 15min at 17,000g and 4°C. After discarding the ethanol, RNA was dissolved in 10µL of RNAse free sterile water (Gibco). cDNA synthesis was performed with the iScript cDNA Synthesis kit (Bio-rad) following the manufacturers’ recommendations. qPCR quantification of CHIKV particles were performed by targeting the *E2* envelope gene coding on CHIKV-positive head, body and saliva samples according to previous FFA titration. *As. taiwanensis* density was estimated by qPCR quantification of the *18S* ribosomal gene in mosquito bodies. For each samples (except saliva), viral gene copies were normalized with copy number of the mosquito housekeeping gene *rps17* coding a ribosomal protein. Specific forward and reverse primers for each gene were listed in **Table S3**. Briefly, quantifications were conducted in a 10µL volume with 1µM of each primer, 1X of Roche SYBR Green I master mix (Roche) and 2.5µL of 1/5^th^ cDNA dilutions. Amplifications were performed on a CFX 1000 thermocycler (Bio-rad) with 5min of denaturation at 95°C, followed by 45 cycles including 10s at 95°C, 30s at 50°C and 10s at 72°C. A fusion step was conducted with 5s at 95°C and a temperature gradient ranging from 65°C to 97°C. For E2 and rps17, 10 times serial dilutions of amplicons were used as a standard after purification with the QIAquick PCR purification kit (Qiagen) following manufacturers recommendations. For the parasite, 10 times serial dilutions of a PCR 2.1-TOPO TA vector (Invitrogen) with an insert of *As. taiwanensis* 18SrDNA was used as a standard.

#### Statistical analysis

Statistical analyses were conducted with the R software v.4.3.3.1. Prevalence data were modeled with a Generalized Linear Mixed Model using a binomial distribution and a logistic link function. Prevalence of the virus (*i.e*. presence – absence) was considered as a response variable while presence of the parasite, body part (including saliva), time post infection and their interactions were used as explanatory variables. Experiment replicates were used as a random variable. Influence of the explanatory variables on the viral prevalence was assessed with a Wald χ^2^ test followed by a *post-hoc* Tukey HSD test with the *car* and *emmeans* R packages respectively. Quantification data were modeled with a Linear Mixed Model (*i.e*. using a normal distribution). Quantification of the virus was considered as a response variable while presence of the parasite, body part (including saliva), time post infection and their interactions were used as explanatory variables. Experiment replicates were used as a random variable. Influence of the explanatory variables on the viral quantification was assessed with a type II ANOVA test followed by a *post-hoc* Tukey HSD test with the *car* and *emmeans* R packages respectively. Correlations between virus and parasite quantification values were conducted with a Spearman test.

**Table S1.** Deviance analyses showing the influence of the time and mosquitoes parasitic status on CHIKV prevalences and densities within each samples types.

| Sample type | Response variable | Explanatory variable | Statistic value (Wald $\chi^2$ / F) | Degree of freedom | p-value |
| --- | --- | --- | --- | --- | --- |
| Body | Proportion of CHIKV positive samples | Time post infection | 0.268 | 2 | 0.875 |
|  |  | Parasitic status | 0.006 | 1 | 0.941 |
|  |  | Time p. i. x Parasitic st. | 0.421 | 2 | 0.810 |
|  | CHIKV copy number among positive samples | Time post infection<sup>†</sup> | 2.316 | 2 | **8.06 x 10<sup>-7</sup>** |
|  |  | Parasitic status | 28.062 | 1 | 0.128 |
|  |  | Time p. i.<sup>†</sup> x Parasitic s | 5.510 | 2 | 0.064 |
| Head | Proportion of CHIKV positive samples | Time post infection | 19.011 | 2 | **7.45 x 10<sup>-5</sup>** |
|  |  | Parasitic status | 8.947 | 1 | **2.78 x 10<sup>-3</sup>** |
|  |  | Time p. i. x Parasitic st. | 2.637 | 2 | 0.268 |
|  | CHIKV copy number among positive samples | Time post infection | 8.985 | 2 | **0.011** |
|  |  | Parasitic status | 3.396 | 1 | 0.065 |
|  |  | Time p. i. x Parasitic st. | 0.195 | 2 | 0.907 |
| Saliva | Proportion of CHIKV positive samples | Time post infection | 3.82 | 2 | 0.148 |
|  |  | Parasitic status | 9.80 | 1 | **1.74 x 10<sup>-3</sup>** |
|  |  | Time p. i. x Parasitic st. | 1.02 | 2 | 0.600 |
|  | CHIKV copy number among positive samples | Time post infection | 0.52 | 2 | 0.772 |
|  |  | Parasitic status | 1.32 | 1 | 0.250 |
|  |  | Time p. i. x Parasitic st. | 3.03 | 2 | 0.230 |

<sup>†</sup>a second order quadratic term was used

**Table S2.** Correlation betwen As. taiwanensis and CHIKV densities.

| Sample type | Time point (day post infection) | p-value | Spearman coefficient (Rs) |
| --- | --- | --- | --- |
| Body | 3 | 0.521 | 0.218 |
|  | 7 | **0.045** | **-0.585** |
|  | 14 | 0.503 | 0.227 |
| Head | 3 | **0.031** | **0.679** |
|  | 7 | 0.059 | -0.616 |
|  | 14 | 0.839 | 0.072 |
| Saliva | 3 | **0.040** | **0.636** |
|  | 7 | 0.377 | 0.296 |
|  | 14 | 0.595 | 0.181 |

**Table S3.**
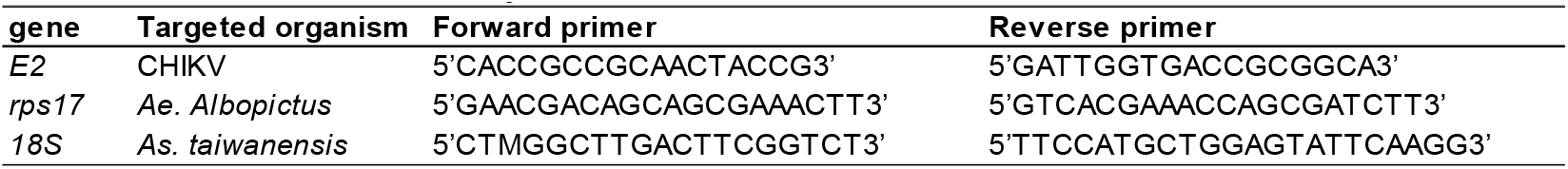
Primers used in the study.

