## Supporting information for "A major entomoparasite interferes with Chikungunya transmission by *Aedes albopictus*"

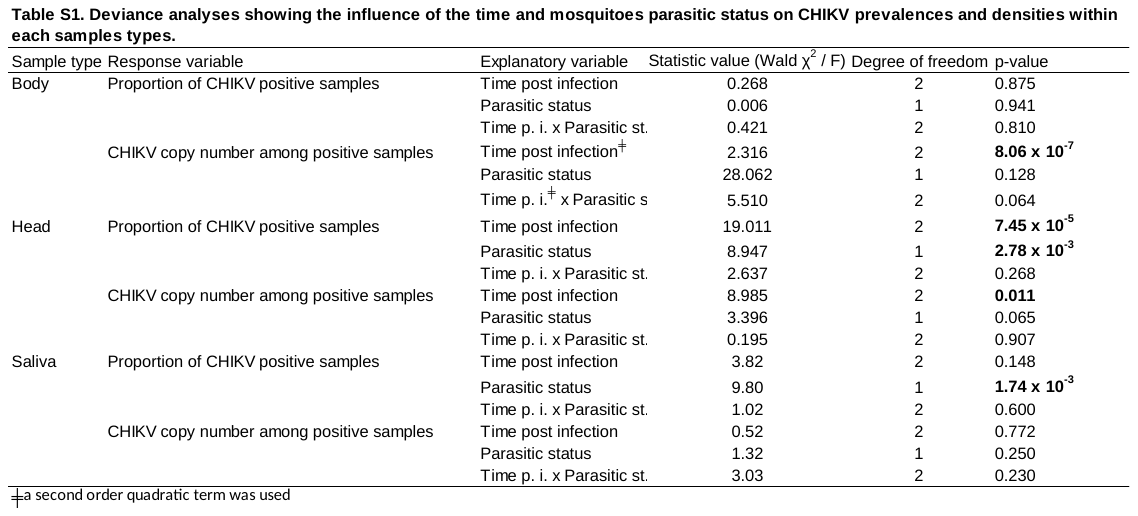

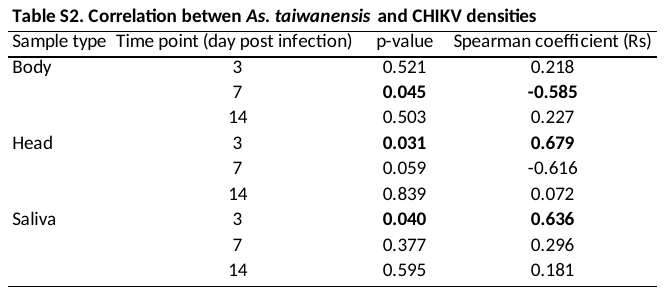

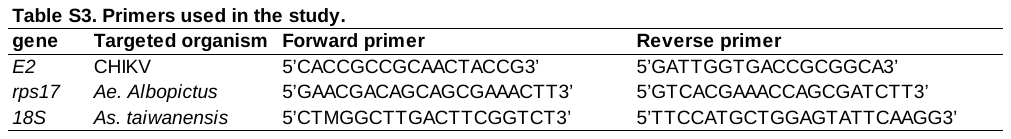
